# Patterns of morphological diversification in the Ramphastoidea reveal the dramatic divergence of toucans from a conserved morphotype

**DOI:** 10.1101/2021.07.29.454288

**Authors:** Krishnapriya Tamma, Anand Krishnan, Sushma Reddy

## Abstract

**Background:** Morphological traits offer insights into an organism’s ecological niche, species interactions, and patterns of community organisation. Pantropical lineages - animals and plants distributed across the three tropical continental regions of Asia, Africa, and America - provide a way to test how different environments and communities influence morphological diversification. Here, we examined a monophyletic group of frugivorous birds, the barbets and toucans (Ramphastoidea), which diversified independently on three continents, to investigate whether clades in each region exhibit similar (phylogenetically constrained) or distinct (ecologically influenced) patterns of morphological diversification.

**Results:** Our results show that despite differences in community dynamics in these regions, lineage accumulation patterns through time on all three continents are broadly similar, putatively due to phylogenetic niche conservatism. We quantified morphological variation in light of phylogenetic relatedness to further reveal that all barbet lineages across continents occupy a conserved region of morphospace after correcting for variation in size. However, in the Neotropics, one lineage, the toucans, have diverged dramatically from typical barbet space and converged toward (yet are distinct from) the trait space occupied by the distantly related hornbills in Asia and Africa. Additionally, we found no link between climatic variables and morphological traits. We conclude that barbets exhibit a conserved morphotype across continents and have diversified by scaling mainly in body size. However, the absence of other large frugivorous birds may have allowed toucans to diversify into a different region of morphospace of increased bill/wing to tail/wing ratios.

**Conclusions:** A combination of photographic and specimen measurements enabled us to demonstrate the presence of a globally conserved barbet morphotype across three tropical continents. By examining different continental lineages of a single monophyletic bird group, we shed light on the contrasting effects of regional ecological factors and phylogenetic constraints on morphological diversification.

## Introduction

Integrating the ecological, evolutionary and biogeographic contexts of lineage diversification has improved our mechanistic understanding of global biodiversity build-up (Jenkins and Ricklefs 2011). The tropics elicit particular interest, because most of the world’s biological diversity is concentrated in these regions (Brown et al 2013). Although tropical and subtropical regions of Asia, Africa and South America have a high degree of endemism, they also share several closely related groups that have diversified independently on each continent. Such pantropically distributed taxa allow us to examine whether diversification proceeds along similar trajectories on different continents, or whether local contingencies such as climate or available niche space shape region-specific diversification patterns (Ricklefs 2004, Phillimore and Price 2008, Burbrink and Pyron 2010, Jønsson et al. 2012, 2015; Tobias et al. 2013, Price et al. 2014, Pigot et al. 2016).

Lineage diversification may be associated with changes in niche occupancy, either by niche expansion or niche packing. Specifically, niche occupancy of a diversifying lineage can change in response to competitive interactions (Brockhurst et al 2007), interactions with predators or pathogens (Ricklefs 2010) or adaptation to novel environments (Lerner et al 2011, Reddy et al. 2012; Tokita et al 2016). An illustrative example is that of the adaptive radiation of Hawaiian honeycreepers driven by the availability of novel habitats in new islands (Lerner et al 2011, Tokita et al 2016). Thus, change in niche occupancy can result in a concomitant change in occupied morphological trait space (or morphospace) for a diversifying lineage. An expectation that follows is that of a positive relationship between morphological disparity and species richness of a lineage (Bell and Barnes 2001, Rabosky et al 2013). Interestingly, studies in passerine birds, the largest avian radiation, uncover evidence of only a weak relationship between morphological diversity and species richness (Ricklefs 2012). This may be a result of phylogenetic constraints on morphology, such as when lineages evolve a large number of morphologically similar species. Examining the morphospace occupied by a clade in the context of phylogenetic relationships between species enables us to examine the relationship between trait diversity and evolutionary history. This, in turn, facilitates insights both into the processes that influence diversification, and how they influence the sorting of species within ecological niche space (Huang et al 2013; Pigot et al. 2020).

When a lineage colonizes different tropical regions, the null expectation is that it occupies congruent regions of morphospace in each region (for example, due to phylogenetic niche conservatism (Moen et al 2013)). However, regional factors including climate and biotic interactions (such as competition stemming from niche overlap) influence patterns of diversity accumulation and trajectories of morphological evolution (Hawkins 2003, Moura et al 2016), thus resulting in some lineages expanding into novel regions of morphospace. One may, therefore, expect that both phylogenetic constraints/inertia and regional factors play contrasting roles in shaping the patterns of niche occupancy within each geographic region. Assuming that regional factors can influence deviation from a conserved morphotype, a key question we aim to address is whether the patterns of morphological space occupied by a single lineage in different regions or continents are similar or distinct.

Here, we studied a pantropically-distributed bird lineage of barbets and toucans (Aves: Piciformes, Ramphastoidea), to compare diversification patterns across three continental regions - tropics of Africa, Neotropics, and Asia. Barbets are largely frugivorous, cavity-nesting birds of tropical and subtropical regions (Goodwin 1964, Short and Horne 2001), which were formerly considered a single family, the Capitonidae. More recent phylogenetic studies (Moyle 2004; add Lanyon & Hall 1994), however, found that toucans were embedded within the barbet clades, and are thus derived from barbets. Barbet lineages on each continent are currently subdivided into different families, the Asian Megalaimidae (2 genera), the African Lybiidae (7 genera), and the Neotropical Capitonidae (2 genera) and Semnornithidae (1 genus). In the Neotropics, barbets are sympatric with the toucans (Ramphastidae, 5 genera), which are generally larger and still retain a mostly frugivorous diet (Remsen 1993). Toucans have been hypothesized to occupy a similar niche in South and Central American forests to that of hornbills in Asia and Africa (Kinnaird and O’Brien 2007, Seki et al. 2010, Van De Ven et al. 2016). This hypothesis is based largely on general appearance and their similar frugivorous diets (McNab 2001), but empirical comparative tests are lacking.

In this paper, we first examined whether barbets in the three continental regions exhibit similar or distinct patterns of lineage diversification. Next, we quantified morphological variation of barbets and toucans to test whether each continental lineage exhibits a conserved or divergent occupancy of morphospace. Further, we examined the toucans to assess whether they occupy a distinct morphospace from other barbets, and if they exhibit convergence with hornbills in Asian and Africa. Differences in morphospace could potentially be driven by regional factors such as climate, which exerts an influence on community composition. Therefore, we tested whether morphological traits on each continent were correlated to patterns in the climate space occupied by each group. In sum, our aim was to conduct a comparative study on the influences of phylogeny and climate on morphological diversification of a widespread tropical group.

## Materials and Methods

### Morphometric data from museum skins

We obtained morphometric measurements from museum study skins of 113 species (out of 127 total) including 33 Asian, 40 African, and 40 Neotropical barbets and toucans held in the collections of the Smithsonian National Museum of Natural History (USNM) in Washington D.C., and the American Museum of Natural History (AMNH) in New York, USA (See Supplementary Data File for total number of skins per species, and museum catalog numbers). Additionally, to test the hypothesis of convergence, we included a representative sample of hornbills from tropical Asia (11 species) and Africa (10 species) for comparison. Using two types of measurements, directly using calipers and digitally via standardized photographs, we took standard measurements of the bill, wing and tail length of specimens (Baldwin 1931). We compared the measurements obtained with calipers and photographs on the same specimens to check whether there were systematic differences. Discrepancy between the two methods was relatively small as a percentage of total length (see Supplementary Material I). For this reason, we felt confident in combining measurements from both methods for further analysis, and using the photographic measurements in those cases where caliper measurements were not available or possible. Example images, and further details on measurements are in the Supplementary Material II, Figure S1.

Using this dataset, we compared morphometric variation between barbet species in two ways: overall size and allometric variation independent of size. For the first, we compared variation in the raw measurements without correcting for size differences between species. Thus, variation in these measurements was heavily correlated to overall size (Ricklefs and Travis 1980). For this same reason, we used the original measurements instead of performing principal components analysis or other dimensionality reduction methods on log-transformed measurements, because linear measurements are all heavily correlated to size and the first component thus typically encompasses almost all the variation.

To examine variation in proportions independent of size, we calculated three ratio measurements: bill:wing, tail:wing and bill:tail ratios for each specimen. To examine the position of toucans relative to barbets in morphospace, we measured misclassification rates by training a classifier to use Linear Discriminant Analysis (LDA) (following Jonsson et al 2015) with five-fold cross validation in MATLAB (Mathworks Inc., Natick, MA, USA) on size-corrected morphological parameters. We implemented this classifier using the Classification Learner app (Chitnis et al 2020), to examine the misclassification rates of each group. Linear Discriminant Analysis requires assignment to pre-existing groups, for which we used current taxonomic groupings (based on the IOC World Bird List accessed from https://www.worldbirdnames.org/new/). Therefore, our six monophyletic groups were the Asian (Megalaimidae), African (Lybiidae), and Neotropical (Capitonidae and Semnornithidae) barbets, toucans (Ramphastidae), and the Asian and African hornbills (Bucerotidae), which we treated as two separate groups to compare individual continental communities. The rationale for this test was that if toucans were convergent with hornbills in morphospace, they should be more likely to be misclassified as a hornbill by LDA rather than as a barbet.

### Comparative phylogenetic analyses

#### i) Building phylogenetic trees

We obtained a distribution of 1000 trees for the Ramphastoidea from the birdtree database (Jetz et al 2012; https://birdtree.org). Using TreeAnnotator (Bouckaert et al 2014), we built a maximum clade credibility consensus tree (ultrametric tree) with 10% burn-in and 0.5 posterior probability limit. Following this, we used pruning functions (e.g., prune.sample) in the picante (Kembel et al 2010) package in R to prune the tree according to the analyses being performed. For instance, when examining variation within the Asian barbets, we pruned the tree to include only those species. For each morphological variable, we used an average value of all specimens of a particular species.

#### ii) Lineage-through-time analysis

We constructed lineage-through-time (LTT) plots, which trace the number of lineages at different time points along a phylogeny, to visualise the overall diversification trends across the three continents. In order to construct these, we used the function ltt.plot from the ape package (Paradis et al 2004) in R (R Core Team 2013), and compared these for the different continental lineages.

#### iii) Comparing morphospace occupancy using phylomorphospace

Phylomorphospace is a method to project a phylogeny onto a morphospace (Sidlauskas 2008), which enables us to examine occupancy of species within lineages in the context of their evolutionary relationships. We constructed phylomorphospace plots using the phytools package in R (Revell 2011), for data on beak length and tail length without size correction, and for the size corrected data based on beak-wing and tail-wing ratios.

#### iv) The role of climate in driving the observed morphological patterns

Using an approach similar to the phylomorphospace, we examined phylogenetic patterns in climate space (characterised by mean annual temperature and mean annual precipitation) occupied by species in each region (henceforth referred to as ‘phyloclimatespace’). To construct this climate space, we first obtained range maps for each species from the IUCN (IUCN 2019), and then determined the mean annual temperature and mean annual precipitation for each range using data derived from WorldClim (Fick and Hijmans 2017), at 1 km resolution. We plotted the ‘phyloclimatespace’ using functions from the phytools package. For purposes of visualization only, the temperature data was scaled (by multiplying by 100) to the same order of magnitude as precipitation. In order to examine whether climate variables were correlated with morphological traits, we performed a phylogenetic generalized least squares regression (PGLS) of each morphological trait against the mean temperature and mean precipitation as predictor variables. For this analysis, we used functions from the R packages ape, geiger (Harmon 2008), nlme (Pinheiro et al 2020) and phytools, and the consensus phylogenetic tree (see above). For each continental lineage, we pruned the tree and the morphological dataset to include only taxa from that continent, and then built three different regression models: one for bill:wing ratio with mean temperature and mean precipitation as predictors, one for tail:wing ratio, and one for bill length alone to examine whether size variation was correlated to climatic space after controlling for phylogeny.

## Results

### Similar patterns of lineage diversification on the three continents

Despite differences in species richness, our lineages-through-time (LTT) analysis of 100 phylogenetic trees revealed that barbet lineages exhibited broadly similar patterns of diversification on the three continents (Figure 1B, C, D). Additionally, LTT plots show that barbet lineages on all three continents exhibit patterns consistent with the ‘late-burst’ scenario, suggesting that the diversification rates on all three continents were initially slow, but exhibited a sharp increase subsequently. Gamma-statistic values (when compared to randomized distributions of 100 trees for each lineage, see Supplementary Material III, Figure S2) support this assertion, as all three lineages exhibit positive values (Neotropics: 1.24, Africa: 0.83, Asia: 2.29). However, only the value for Asia was statistically significant.

**Figure 1:**
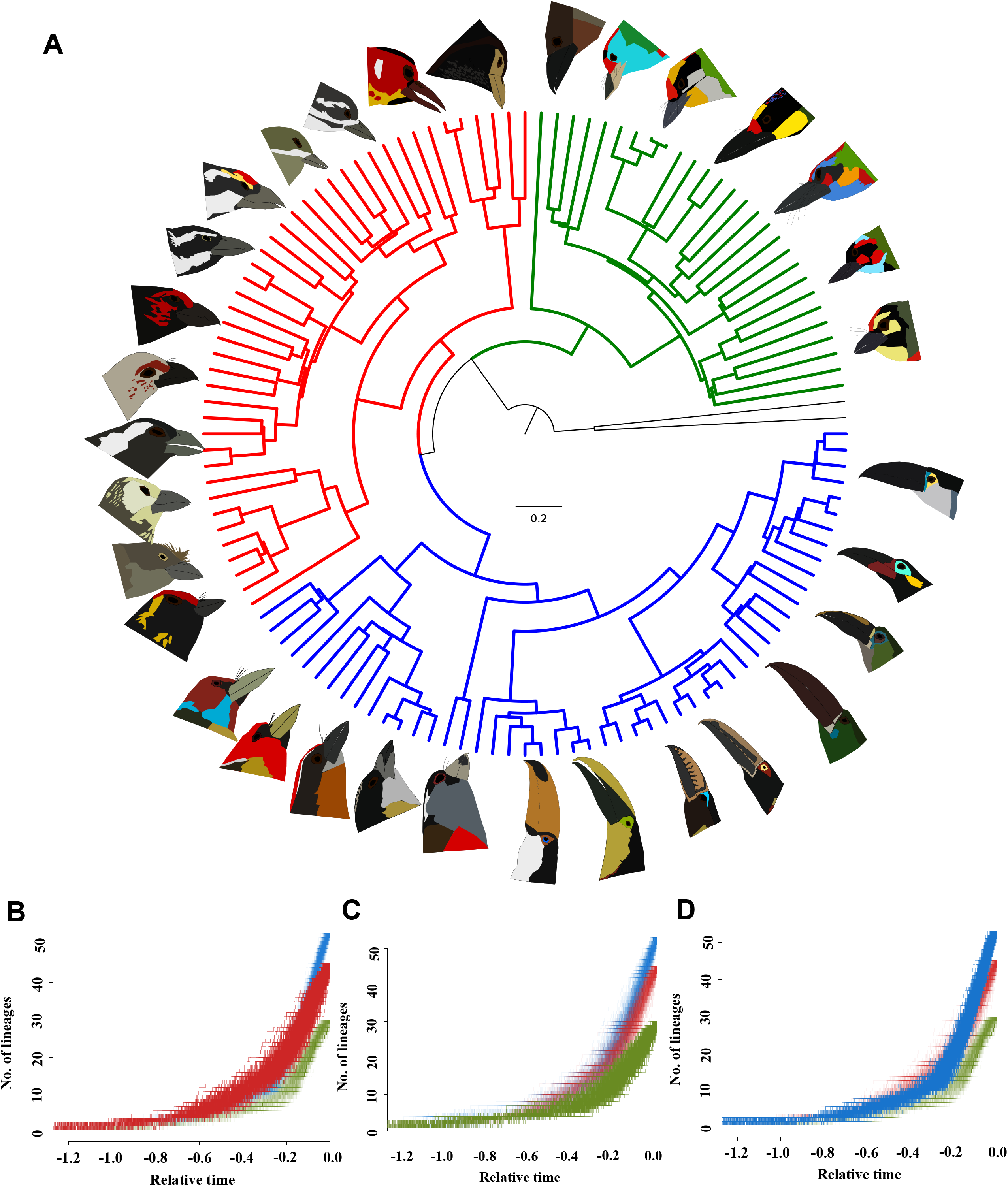
(A)Phylogenetic relationships of barbets and toucans showing distinct continental radiations in Asia (green), Africa (red) and the Neotropics (blue). The tree represents a maximum clade credibility consensus tree of the barbet/toucan clade from birdtree.org. Drawings by A. Krishnan. Lineages-through-time (LTT) plots for each continental clade (sampled across 1000 possible trees) are shown with Africa (B), Asia (C), and Neotropical (D) regions highlighted on top and the other two in background for comparison. Barbets on these three continents exhibit similar patterns of lineage accumulation.

### Morphological diversification patterns differ across continents

From the phylomorphospace for both raw (tail length and beak length) (Figure 2A), and size-corrected (bill-wing ratio and tail-wing ratio) measurements (Figure 2B), we found that barbets in the Neotropics, Africa and Asia occupy similar morphospace, with toucans distinct from all other lineages. For the size-corrected measurements, 3D phylomorphospace using all three ratios did not differ from a 2D phylomorphospace (Supplementary Material IV, Figure S3). Therefore, for easy comparison of size-corrected and raw measurements, we show the bill:wing and tail:wing ratios plotted against each other.

**Figure 2:**
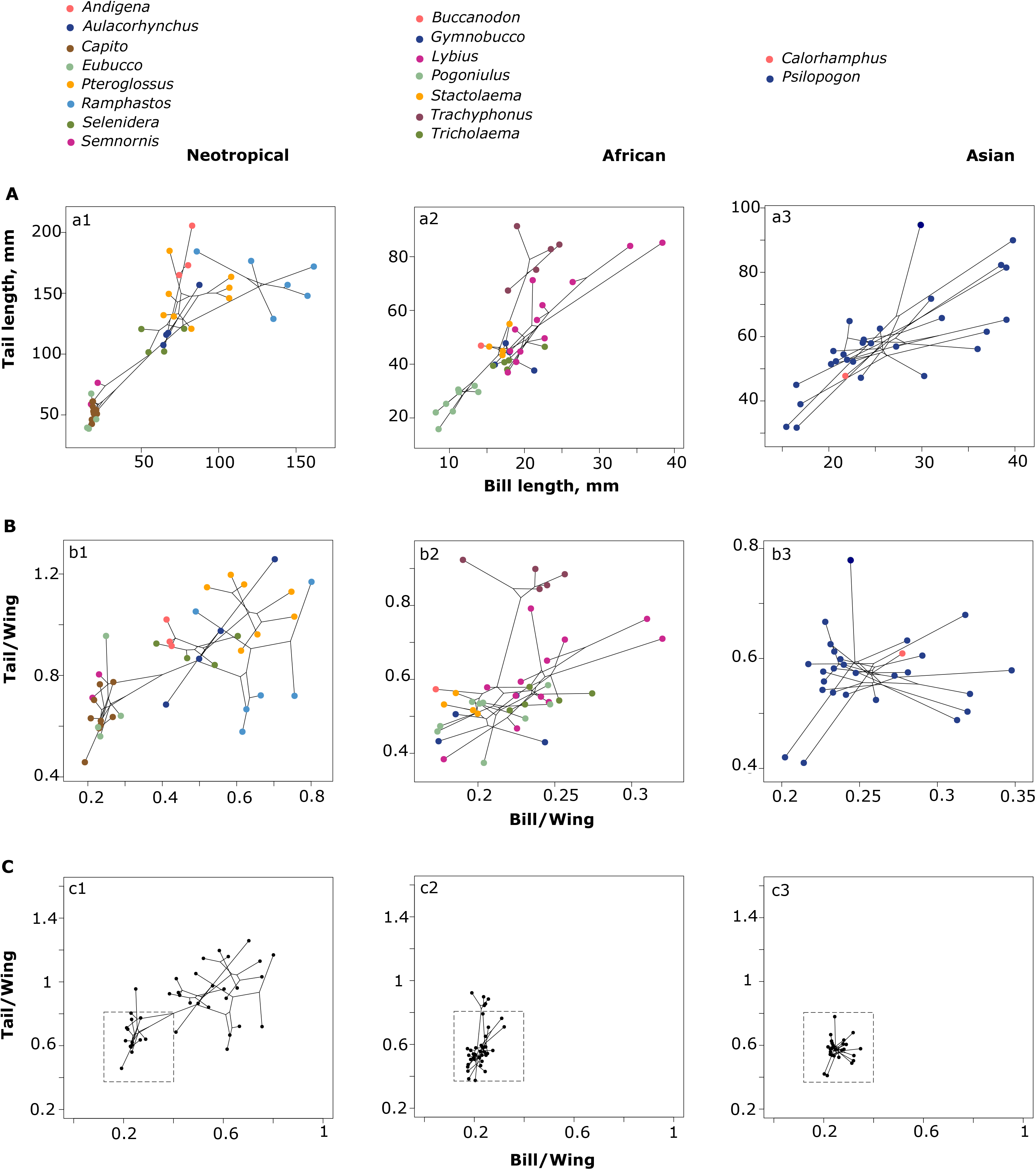
A) Phylomorphospace analysis of barbets and toucans showing size variation across groups. Bill length in relation to tail length for Neotropical (left), African (center) and Asian (right) lineages. Each point represents a species, with genera color-coded according to the key. B, C) Phylomorphospace of size-corrected measurements where bill and tail lengths are standardized using wing length; (B) shows details of continental variation while C) shows the same plots but scaled to similar axes for comparison across continental groups. The dashed box represents the approximate extent of the ‘conserved barbet’ morphospace occupancy. African barbets generally occupy the same conserved region of morphospace as Asian barbets, except for the genus *Trachyphonus*. In the Neotropics, toucans have undergone a dramatic expansion into a novel region of morphospace, a pattern evident even after correcting for the larger size of toucans.

For all three continents, we observed correlated variation in bill and tail lengths in the Ramphastoidea (Figure 2A). However, when size-corrected ratios were used, this correlation was less evident (Figure 2B). Toucans form a separate cluster from other Ramphastoidea even after size correction, implying that they exhibit considerable divergence in body proportions as well (Figure 2B). Euclidean distances between the centroids of each group in size-corrected morphospace support the assertion that barbets (i.e. excluding toucans) are very close to each other in morphospace (Asian-African barbets: 0.0375, Asian-Neotropical barbets: 0.0988, African:Neotropical: 0.0816). All three barbet groups were highly distinct from the toucans (Euclidean distances between the centroids in size-corrected morphospace: Toucans-Asian barbets: 0.5024, Toucans: African barbets: 0.5134, Toucans: Neotropical barbets: 0.4491). We defined a ‘constrained barbet morphospace’ using the morphospace occupied by the Asian barbets (Megalaimidae), because this clade has the lowest species diversity and morphological disparity (dashed-line box in Figure 2C). When we map this ‘constrained barbet morphospace’ onto the other continents, we found that data points from the African (Lybiidae, with the exception of *Trachyphonus)*, and the Neotropical (Capitonidae and Semnornithidae) species remained within the area delimited by the Asian clade. This supports the idea of a constrained ‘barbet morphotype’ across continents, except the toucans, which deviate substantially from this morphospace. There was no obvious additional clustering within barbets (Figure 2C).

### Toucans are convergent with hornbill morphospace

The results of the morphospace analysis demonstrate that barbets across all three continents occupy a clustered region of morphospace when corrected for size (Figure 2C), whereas toucans (which are nested within the clade of Neotropical barbets) occupy a distinct region of morphospace. Toucans are distinct from all other barbets (see above), and their morphospace is closer to (although still distinct from) the Asian and African hornbills (Figure 3A and B). Asian and African barbets are very distinct from hornbills on their respective continents (Euclidean distances of 0.3363 and 0.3998 respectively). Finally, toucans are much closer to Asian (0.1829) and African (0.2152) hornbills in morphospace than both are to any of the barbet groups.

**Figure 3:**
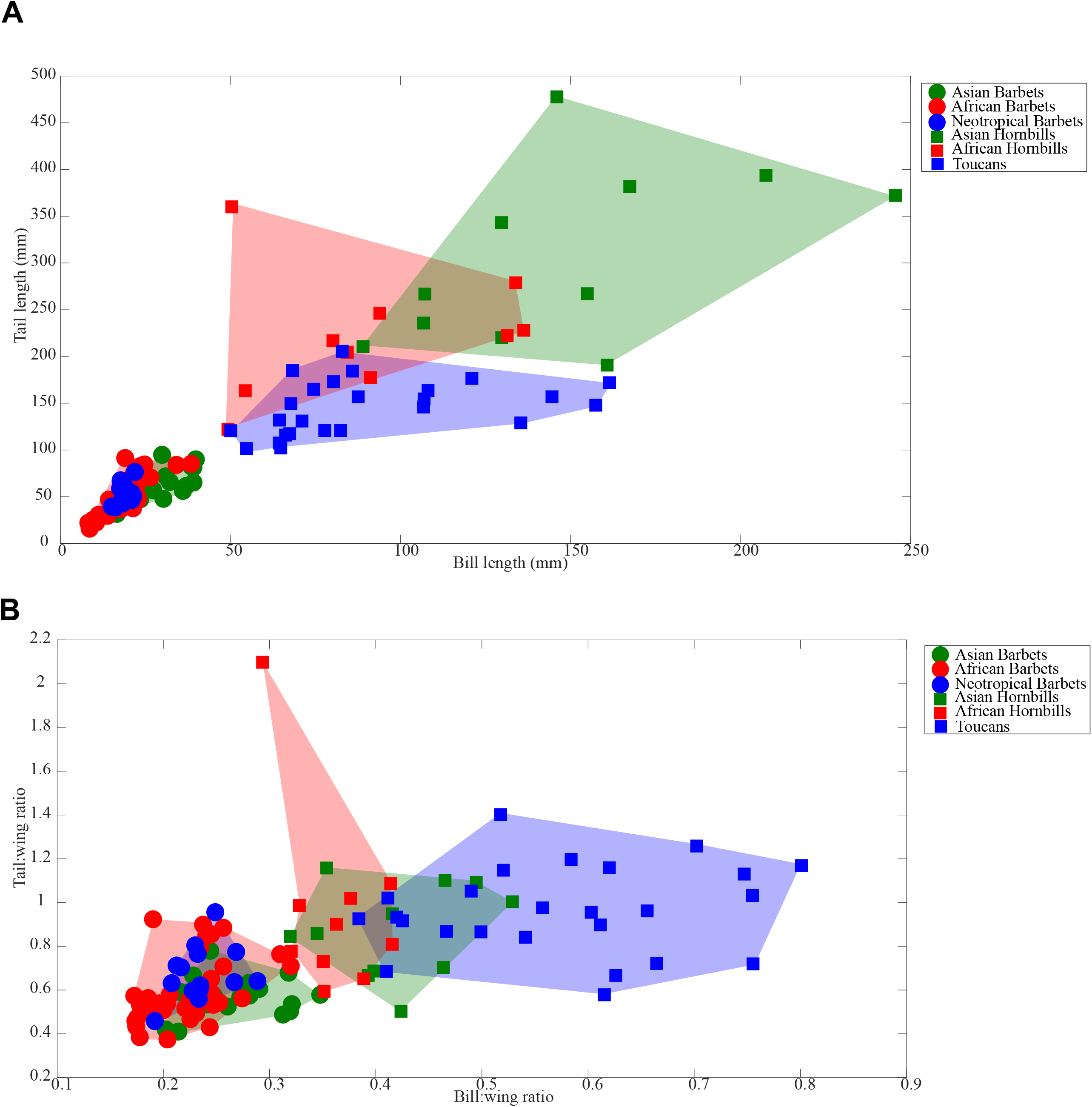
Comparative morphospace of barbets, toucans, and hornbills. (A) Bill-length versus tail-length (in mm) for barbets, toucans and hornbills. Groups are colored by continent; circles indicate barbets and squares are toucans and hornbills. (B) Size-corrected morphospace of bill:wing ratio against the tail:wing ratio. Barbets across continents are clustered within an overlapping region of morphospace, whereas toucans form a distinct cluster that overlaps hornbill morphospace.

Classification using Linear Discriminant Analysis in size-corrected morphospace further supports our assertion that toucans are closer to hornbills in morphospace. Barbets from any one continent were most frequently misclassified as another of the two barbet groups (Table 1), but were never misclassified as a toucan. This indicates heavy overlap of barbet groups in morphospace, but distinctness from toucans and hornbills. Likewise, the toucans were never misclassified as a barbet, but were misclassified as hornbills a total of 24% of the time. This provides quantitative support for toucans having converged toward hornbill morphospace and away from other barbet groups, although they do not completely overlap with hornbill morphospace.

**Table 1:**
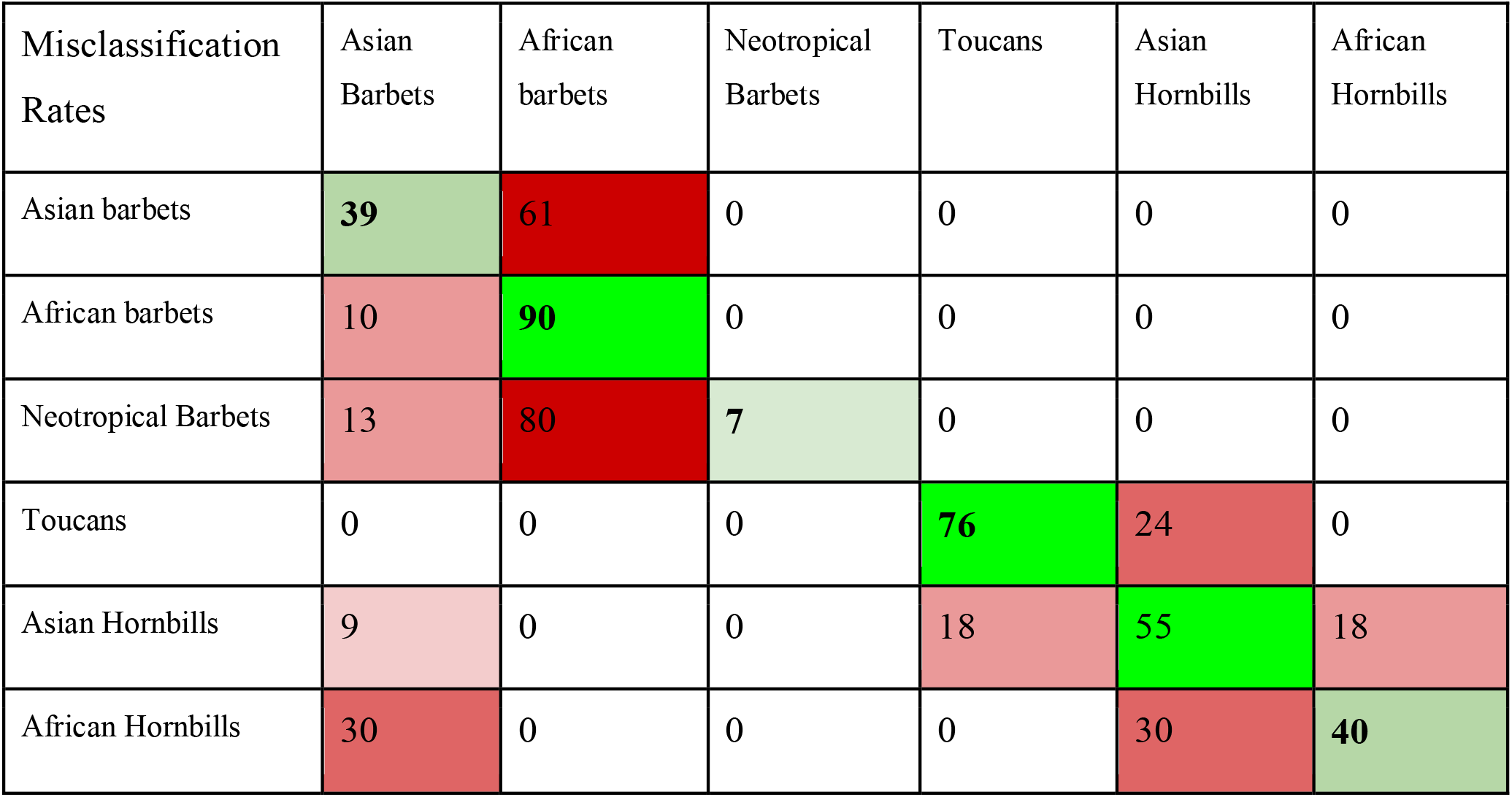
Percentage misclassification rates using Linear Discriminant Analysis on size-corrected morphological variables. The closer two groups are in morphospace, the more likely they are to be misclassified as each other. Bold numbers along the diagonal (green squares) indicate the correct classification rates for each group, with brighter green shades indicating higher rates. Red squares indicate misclassification rates, with brighter red shades indicating a higher rate of misclassification. Note that barbets were often misclassified as other barbets from different regions but were never misclassified as a toucan. Toucans, on the other hand, were misclassified only as hornbills and never as a barbet.

| Misclassification Rates | Asian Barbets | African barbets | Neotropical Barbets | Toucans | Asian Hornbills | African Hornbills |
| --- | --- | --- | --- | --- | --- | --- |
| Asian barbets | **39** | 61 | 0 | 0 | 0 | 0 |
| African barbets | 10 | **90** | 0 | 0 | 0 | 0 |
| Neotropical Barbets | 13 | 80 | **7** | 0 | 0 | 0 |
| Toucans | 0 | 0 | 0 | **76** | 24 | 0 |
| Asian Hornbills | 9 | 0 | 0 | 18 | **55** | 18 |
| African Hornbills | 30 | 0 | 0 | 0 | 30 | **40** |

### African barbets generally occupy a drier climate space

The ‘phyloclimatespace’ for the three continental barbet radiations exhibited no clearly defined pattern within each continent (Figure 4). This is especially evident for species in the Neotropics, where we do not recover the two distinct clusters observed in the phylomorphospace (Figure 2, a1). Neotropical and Asian genera occupy a comparable climate space, whereas African barbets generally occupy a drier climate space, characterised by lower mean precipitation (and are displaced to the left of the other two lineages in the ‘phyloclimatespace’, see Figure 4). The minimum and maximum precipitation values (species means of the mean annual precipitation, in mm) for African, Asian and Neotropical groups are 1641.419 - 420.4235 (max - min), 3324.164 - 1002.519, and 3834.843 - 1058.088 respectively., showing that the African species occupy a drier regime. However, across all continents, there are no observable clade-specific patterns, with the exception of the forest-restricted African genus *Gymnobucco*, which occupies wetter habitats compared to other African barbets.

**Figure 4:**
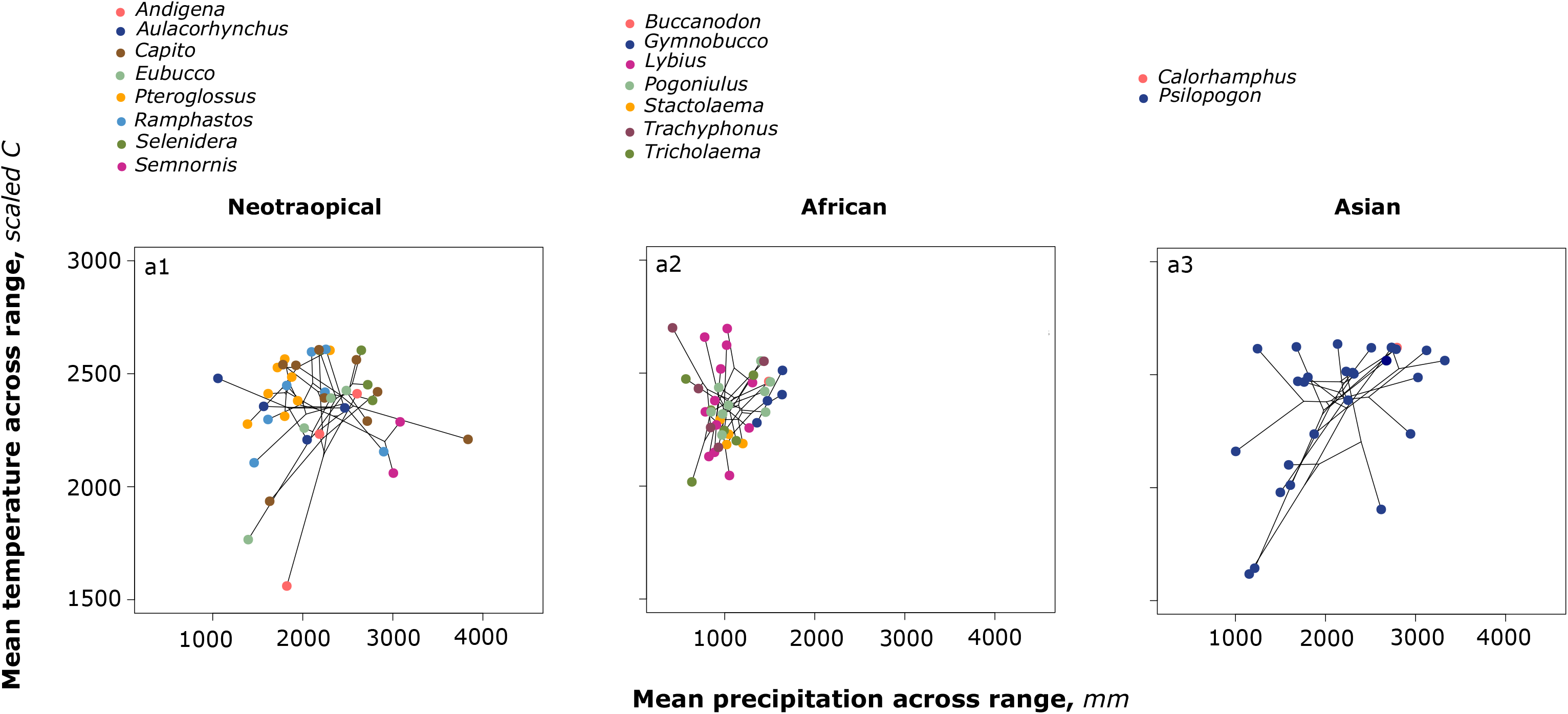
“Phyloclimatespace” plots for each continental radiation, South America (a1), Africa (a2) and Asia (a3). Genera are color-coded in the same way as Figure 2. These plots ordinate mean precipitation against mean temperature for each species range (temperature is scaled to the same range of values as precipitation), obtained from the WorldCLIM dataset using species ranges from the IUCN Red List. African species occupy a distinct climate space from Asian or Neotropical species.

The phyloclimatespace results were consistent with the results of the PGLS analyses summarized in Table 2. After controlling for the effects of phylogenetic non-independence, we observed no significant relationship between temperature, precipitation and morphological variables for the African barbets. For the Asian barbets, bill length exhibited a significant (P<0.05) relationship with mean precipitation, but all other relationships were not statistically significant. For the Neotropical barbets and toucans, only tail:wing ratio is significantly correlated with mean precipitation. Thus, although two out of 27 correlations between climatic and morphological traits were statistically significant, we did not uncover any broad patterns of morphological diversification being correlated to diversification in temperature-precipitation climate space.

**Table 2:**
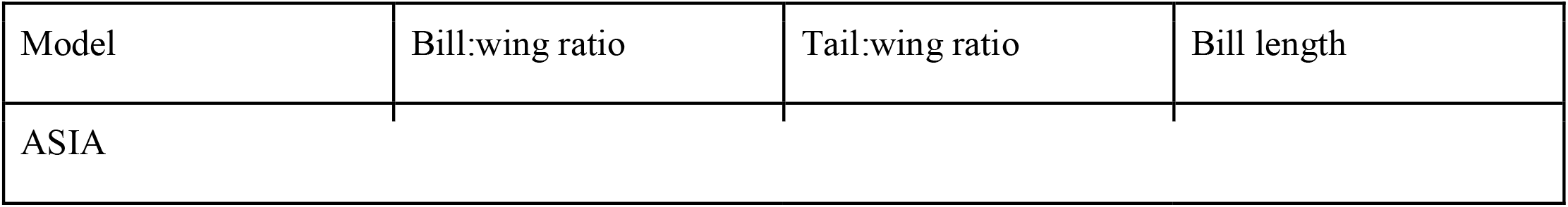
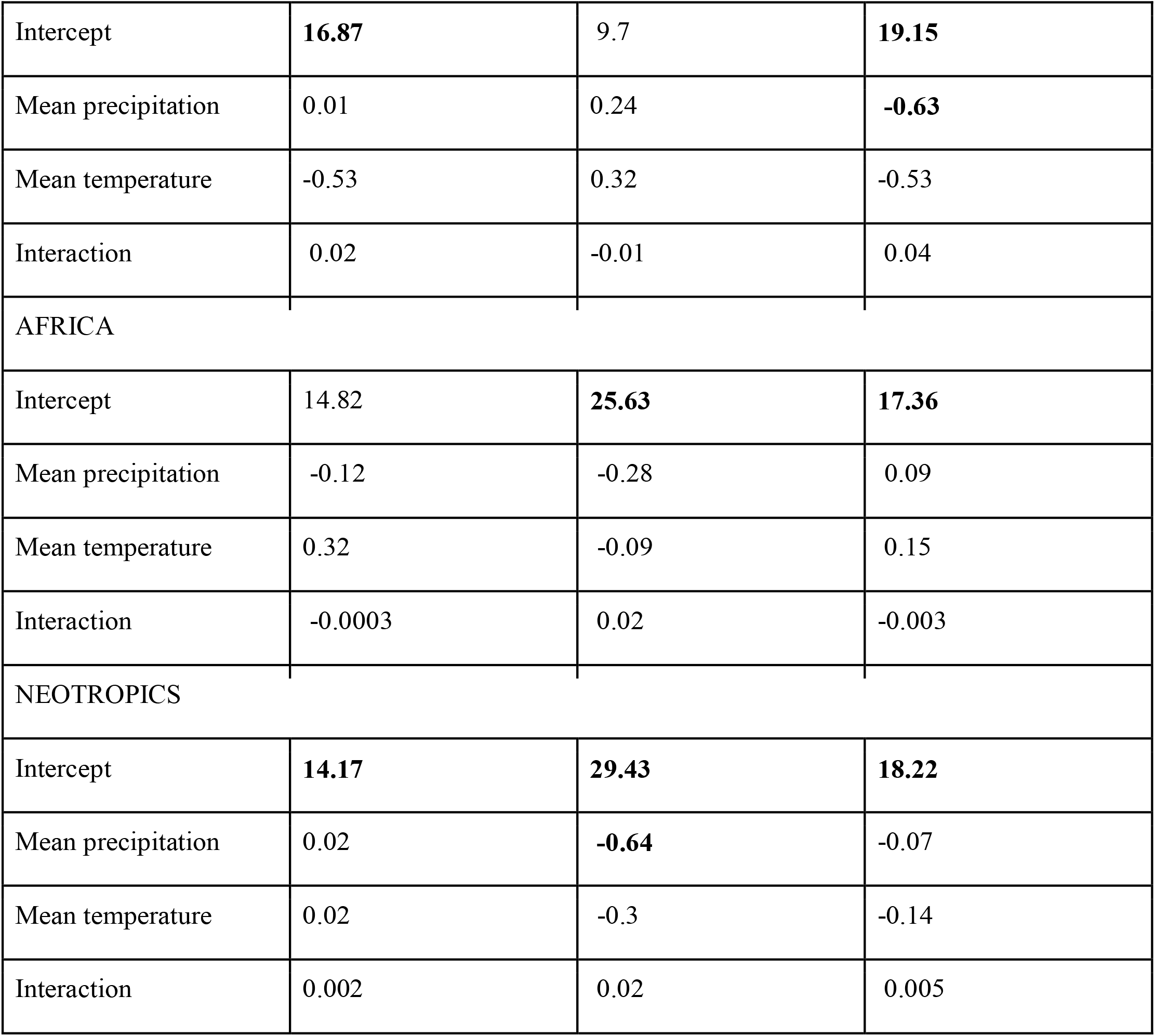
Results of phylogenetic generalized least squares models comparing climatic and morphological variables. Values shown represent the coefficients, with statistically significant results (P<0.05, for comparison of independent and dependent variables) in bold.

## Discussion

In this study, we investigated patterns of diversification and morphospace occupancy in the pantropically distributed barbets and toucans (Ramphastoidea) at continental scales, to understand how regional factors influenced the evolutionary trajectories of related bird groups occupying different areas. Patterns of lineage accumulation in the Ramphastoidea show similarities between the three continents. Using morphological data collected from museum specimens, we recovered a constrained ‘barbet morphotype’ that occupies the same morphospace on all tropical continents. Additionally, in the Neotropics, there was a dramatic expansion into a novel morphospace, driven entirely by the diversification of toucan lineages. Finally, we demonstrated using quantitative data that toucans are not only highly distinct from all other barbets, but also converge upon the morphospace occupied by hornbills in Asia and Africa.

Our morphological dataset used museum specimens in a somewhat novel approach of combining measurements taken directly on specimens along with those from photographs. By cross-verifying these measurements, we found them suitable to merge. We hope that photographs will provide a more accessible source of morphometric data, particularly in the era of digitized museum collections. Because of the prohibitive cost of traveling to museums, particularly for researchers in developing countries, photo datasets can provide a powerful alternative to conventional morphometric methods.

A conserved barbet morphology across the globe suggests the existence of constraints to the evolution of morphospace in barbets. This is supported by comparative morphometric studies in other taxa: for example, studies on passerine birds demonstrated that lineages on four continents have evolved in a congruent fashion, occupying similar morphospaces despite independent evolutionary trajectories (Ricklefs 2012, Jonsson et al 2015). This tendency to conserve aspects of their ancestral niche suggests that morphospace of species within a diversifying lineage is potentially shaped by evolutionary constraints. The radiation of toucans in the Neotropics is consistent with a scenario of release from such evolutionary constraints. We found support for this assertion from our preliminary disparity-through-time (DTT) (see Supplementary Material V, Figure S4), with Africa and the Neotropics showing patterns consistent with late-burst, suggesting within-clade diversifications. Such a tendency also points to the oft-ignored role of non-adaptive radiations within lineages, where phylogenetic diversification is not accompanied by morphological diversification (Reaney et al 2018).

Under certain ecological conditions such as the absence of competitors (ecological opportunity), pathogens (parasites) and predators, lineages may undergo rapid diversification and expansion in morphospace (Lovette et al. 2002, Harmon et al. 2003, Burbrink and Pyron 2010, Yoder et al. 2010, Ricklefs 2010, Jønsson et al. 2012). Phylogenetic analyses suggest that crown hornbills and barbets likely originated at roughly similar times (Prum et al. 2015, Claramunt and Cracraft 2015), and it is thus possible that the presence of hornbills or other frugivores has constrained the morphospace of barbets in Africa and Asia. In the Neotropics, however, where hornbills are absent, one lineage of barbets (the toucans) has undergone a dramatic shift toward hornbill morphospace, evolving proportionately longer bills as well as longer tails.

Although climate is an important driver of diversification in the tropics (Castro-Insua et al. 2018), we do not observe any discernible phylogenetic trends in the occupancy of climate space, and only two out of 27 correlations between climatic and morphological variables were statistically significant. Primarily, we find that African barbets occupy generally drier climate spaces than those on the other continents; in spite of this, however, their morphospace is congruent with the Asian and Neotropical barbets. This is consistent with African barbets generally inhabiting savanna and scrub habitats, as compared to barbets on other continents, which generally inhabit wetter forest habitats (Short and Horne 2001). The climate trait space of African barbets may result from prehistoric climate transitions between wet and dry habitats that have shaped the modern biogeography of the African continent (Tolley et al 2008, Ivory et al 2018). However, this does highlight just how robust the conserved morphology (and thus niche) of barbets is across a number of different habitat types. Further, this is consistent with our earlier explanation of ecological opportunity, rather than changes in climate, driving the dramatic divergence of toucans from Neotropical barbets.

## Conclusions

To conclude, our results were consistent with the presence of a globally conserved barbet morphotype across three tropical continents (Africa, Asia and South America). We present preliminary evidence that ecological opportunity (i.e. the absence of hornbills in South America) putatively resulted in a disjunct expansion to novel morphospace during the diversification of toucans from a barbet ancestor. Our results are a key step in understanding whether lineages diversifying in different regions show conserved evolutionary trajectories, and thus have implications in understanding how the enormous amount of diversity in the tropics evolved.

## Acknowledgments

We thank Lamiya S for taking independent measurements of the barbet specimens which were used to ensure that our measurements (both calipers and direct) were accurate. We thank Chris Milensky, Brian Schmidt, Christina Gebhard and Helen James from the Smithsonian National Museum of Natural History, and Paul Sweet and Lydia Garetano from AMNH, for providing requisite permissions to access samples and support.

## Author contributions

KT and AK conceived of the study, and all authors conceived the methodology and analyses; AK photographed specimens and made caliper measurements; KT photographed additional specimens. KT and AK obtained measurements of specimens from photos; KT and AK analysed the data; AK, KT and SR contributed to data interpretation and writing. All authors read and approved the final manuscript.

