## Supplementary Material for "Patterns of morphological diversification in the Ramphastoidea reveal the dramatic divergence of toucans from a conserved morphotype"

### I. Additional Information on Measurements

We took morphological measurements in two ways: 1- directly on specimens using Vernier calipers ; 2- from standardized photographs of the specimens. To reduce the time needed for the collection of data from multiple specimens, we used this combined approach. We strived to maintain the accuracy of morphometric measurements by consistently photographing specimens in the same orientations : (1) lateral view of the body with culmen parallel to the lens to allow consistent measurements of bill and wing chord length, and (2) dorsal view of whole body to measure tail length. Using ImageJ (NIH, Bethesda, MD, USA) and the scale bar as a reference, we made three measurements that corresponded to direct measurements on specimens using calipers: (1) the length of the bill at culmen in a straight line from the base of the bill (where feathers ended) to tip (corresponding to Fig 3 in Baldwin 1931), (2) length of wing chord from the shoulder to the wingtip in a straight line (Fig 100-101 in Baldwin 1931), (3) tail length in a straight line from the base of the tail feathers to the tip of the longest feather (Fig 120 in Baldwin 1931).

From a previous study (Krishnan and Tamma 2016), we had both caliper and photographic measurements from 27 Asian barbet specimens. When comparing each pairwise bill length measurement of the same individual, the average discrepancy between measurements was roughly 2.5mm, and the median was 1.9mm, whereas variation between individuals of a species, measured using the same method, may be higher than 5mm (Krishnan and Tamma 2016). For 7 specimens of Neotropical Barbet, we also quantified the individual variation in measurements from photographs as a measure of digitization error, by having a second person repeat these measurements. The average discrepancy in bill length using this method was 0.8mm (the specimens had bill lengths between 13-16mm), for tail length was 4mm (the tails of these specimens were between 39-61mm), and for wing length was 8mm (the wings of the specimens measured between 59-97mm).

### II. Measurements of morphological trait data from museum specimens

Example photographs showing, for illustrative purposes, how we measured bill, tail and wing lengths. Specimens were photographed in three orientations, and the measurements were made from whichever photograph was clearest, and where bending of the specimen would not influence the measurement taken. Bill (culmen length) represented the linear distance from the base of the maxilla to the tip, wing chord length represents the length of a straight line from the shoulder to the wingtip, and tail length was measured from the tail base to the tip of the longest tail feather as shown in the examples below

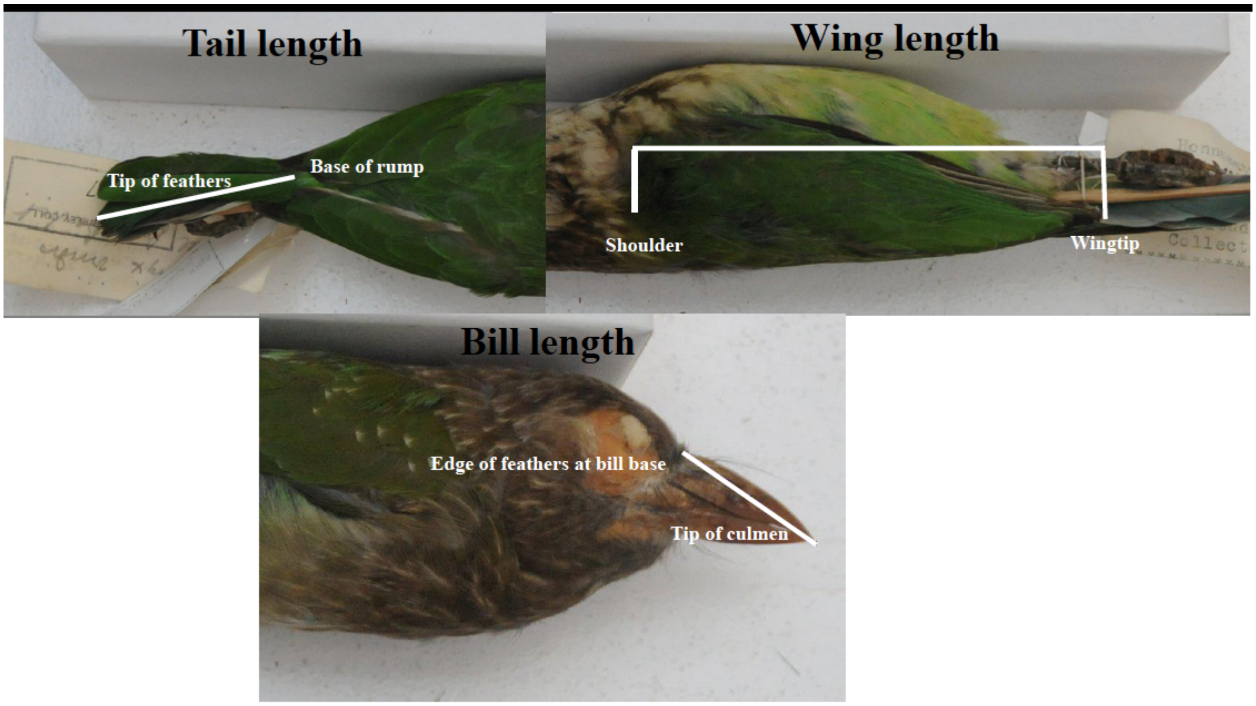


Figure S1: Images of barbet specimens highlighting the measurements taken on each.

### III. Gamma values for LTT plots

The lineage-through-time (LTT) plots were constructed in R using functions from the package phytools. The significance of the gamma statistic was estimated using the MCCR test with 100 simulations. The observed gamma value is marked on a plot of the frequency distributions of the simulated gamma values. The gamma value for Asia is statistically significant.

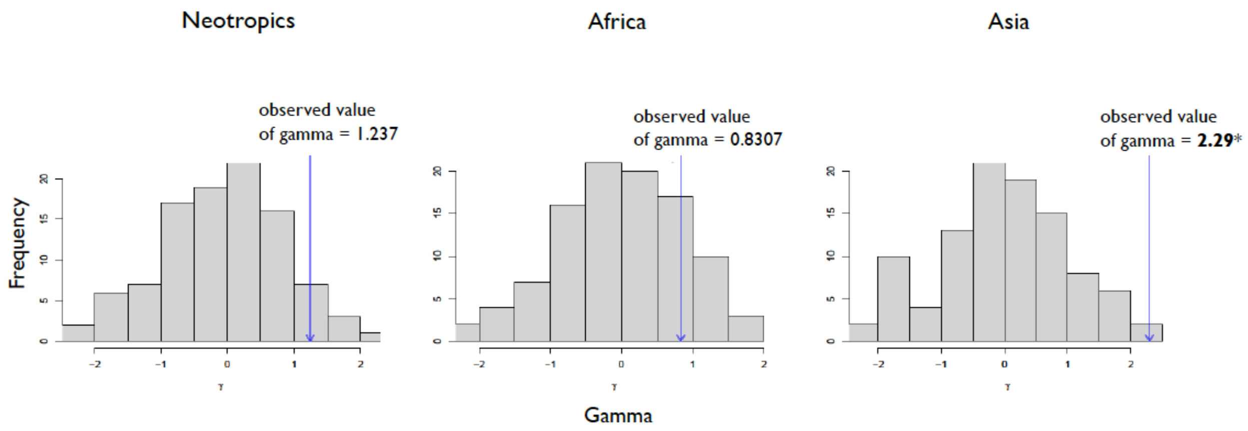


Figure S2: A plot of the frequency distribution of the 100 simulated gamma values for each continent. The observed gamma values are indicated in the plots.

### IV. Phylomorphospace with Bill/Tail, bill/wing and tail/wing ratios

We constructed 3D phylomorphospaces with bill:tail, bill:wing and tail:wing ratios. The patterns are consistent with what we observe for the 2D phylomorphospace for size corrected data. This suggests that the bill:wing and tail:wing ratios are capturing the patterns in the morphological data for these groups.

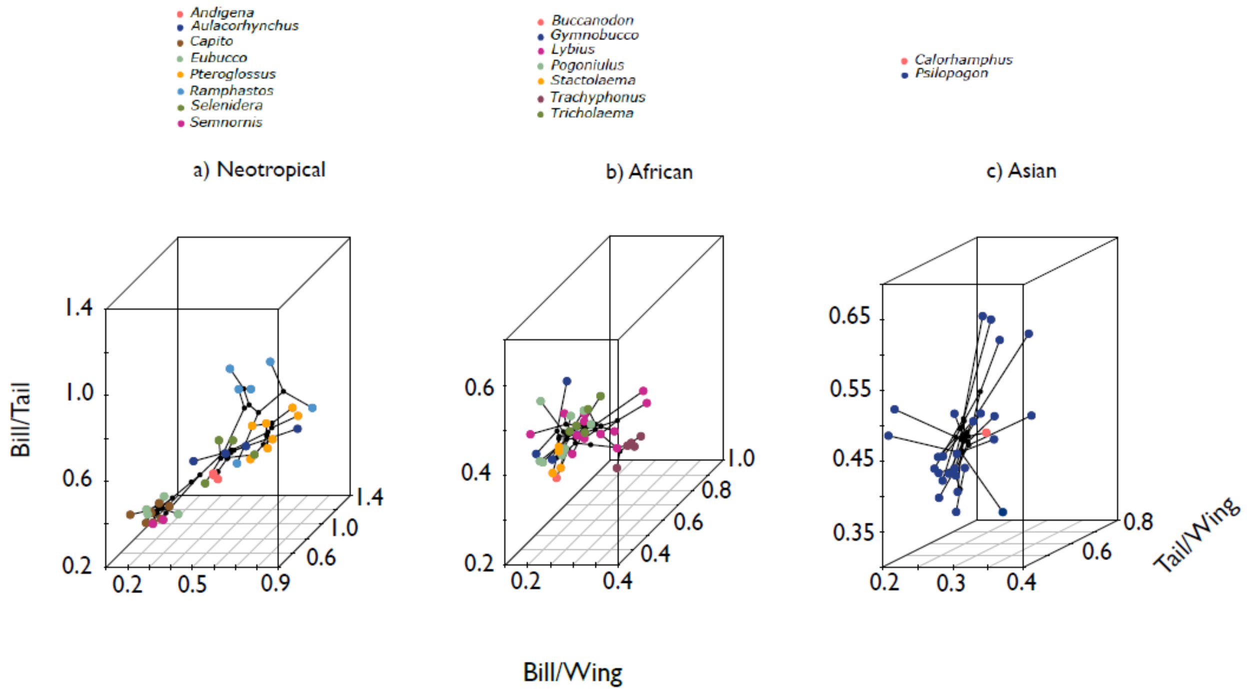


Figure S3: The 3D phylomorphospace for the three continents based on bill:tail, bill:wing and tail:wing ratios

### V. Disparity-through-time (DTT) analyses of morphological diversification

A disparity through time (DTT) plot represents the mean disparity (morphological variation) of each subtree (or clade) in comparison to the disparity of the whole lineage. DTT plots allow us to examine whether disparity in the morphological trait under consideration is mostly accumulated within or across subclades. Functions from the geiger package in R were used to calculate disparity and construct disparity-through-time (DTT) plots. We combined the consensus tree obtained using TreeAnnotator as described above with the morphological data for beak-wing ratio and tail-wing ratios to calculate disparity and construct the DTT plots. The DTT method first calculates the mean relative disparity for all the subclades at different points in time. This observed mean relative disparity is then compared to the median relative disparity calculated from the null patterns of trait evolution, obtained via 10,000 simulations of the trait under Brownian motion. This comparison can be quantified using the Morphological Disparity Index (MDI). MDI is calculated as the area between the two curves - that is, the mean observed disparity and the median of the null disparity values (from 10000 simulations). Values of MDI above 0 suggest that most of the morphological disparity is partitioned within subclades, while values of MDI below 0 suggest that most of the morphological disparity is partitioned between subclades. For each region, we obtained a distribution of MDI estimates from 100 runs of the dtt.plot function, and computed the mean and the standard error.

Disparity-through-time (DTT) analyses support the assertion of continent specific diversification processes within barbet lineages. Observed subclade disparity for African barbets is consistently lower than simulated values, whereas for Asian barbets subclade disparity is consistently higher. Neotropical barbets exhibit higher observed subclade disparity than expected earlier in their diversification, and a decrease in observed disparity over time to a lower value than the simulation.

MDI values are most negative and statistically significantly different from the median simulated disparity (p-value = 0.025; mean = -0.272, se = 0.000151) in African barbets, which indicates that most of the morphological disparity is partitioned between subclades (and thus the presence of clade-specific morphotypes). For Neotropical clades, the MDI value is weakly negative ( p-value = 0.564; mean = -0.048, se = 0.000132). The MDI value for Asian barbets is positive (p-value = 0.888; mean = 0.344, se = 0.000089) but not significantly different from the median simulated disparity, suggesting that diversity is generally partitioned within subclades.

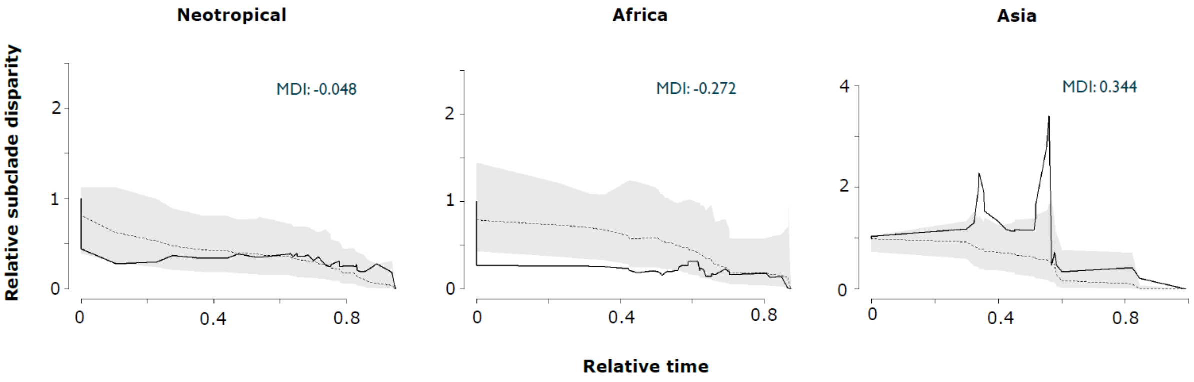


Figure S4: Disparity-through-time (DTT) plots for each continental radiation of barbets.

(although the value of this index is not statistically significant for Neotropical Barbets; we did not construct these graphs for Asian barbets owing to low taxonomic sample size, and as such report DTT plots here only for preliminary comparison)
